# Loss of α7 nicotinic acetylcholine receptors in GABAergic interneurons causes sex-dependent impairments in postnatal neurogenesis and cognitive and social behavior

**DOI:** 10.1101/2020.03.16.994111

**Authors:** Samir A. Nacer, Simone Otto, Ayland C. Letsinger, Jemma Strauss DeFilipp, Viktoriya D. Nikolova, Natallia V. Riddick, Korey D. Stevanovic, Jesse D. Cushman, Jerrel L. Yakel

**Author notes:** Corresponding Author: Jerrel L. Yakel, Neurobiology Laboratory, National Institute of Environmental Health Sciences, National Institute of Health, Department of Health and Human Services, Research Triangle Park, P.O. box 12233, Mail Drop F2-08, North Carolina, 27709, USA.

## Abstract

Neural stem cells within the subgranular zone of the dentate gyrus (DG) generate new neurons that form the granule cell layer during embryonic development and continue to generate new neurons throughout life. The maturation process of newly generated granule cells is modulated by nicotinic acetylcholine receptors (nAChRs), which have been shown to play a role in cell survival, signal modulation, dendritic integration, and memory formation. Disrupted nAChR signaling has been implicated in neuropsychiatric and neurodegenerative disorders, potentially via alterations in DG neurogenesis. GABAergic interneurons are known to express nAChRs, particularly the α7 subunit, and have been shown to shape development, integration, and circuit reorganization of DG granule cells. Therefore, we examined the effects of conditional deletion of α7 nAChRs in GABAergic interneurons on measures of postnatal neurogenesis and behavioral outcomes. Loss of α7 nAChRs resulted in a decrease of postnatal granule cells, as indicated by reduced GFAP+ cells in the DG, specifically in male mice, as well as sex-dependent changes in several behaviors, including social recognition, object investigation, and spatial learning. Overall, these findings suggest α7 nAChRs expressed in GABAergic interneurons play an important role in regulating postnatal neurogenesis and behavior in a sex-dependent manner. This provides important insight into the mechanisms by which cholinergic dysfunction contributes to the cognitive and behavioral changes associated with neurodevelopmental and neurodegenerative disorders.

## Introduction

α7 nicotinic acetylcholine receptors (nAChRs) are highly expressed within excitatory and GABAergic interneurons in the hippocampus where they play an important role in supporting normal cognitive function via a variety of proposed mechanisms (Gotti et al. 2006; Haam and Yakel 2017). Aberrant α7 nAChR signaling has been implicated in neuropsychiatric and neurodegenerative disorders, such as schizophrenia and Alzheimer’s disease. Single nucleotide polymorphisms resulting in overexpression of a dominant negative version of the CHRNA7 gene (CHRNA7A) as well as reduced expression of α7 nAChRs have been found in the brains of schizophrenic patients (Stevens et al. 2014; Kunii et al. 2015). Furthermore, positive allosteric modulators of α7 nAChRs have been investigated as a potential treatment for a variety of disorders, including Alzheimer’s disease in which the cholinergic system diminishes (Bertrand et al. 2015; Dineley et al. 2015; Ma and Qian 2019). Thus, understanding the processes of α7 nAChRs signaling in the hippocampus is critical to understanding normal brain function and developing treatments for neuropsychiatric and neurodegenerative disorders.

Neural stem cells (NSCs) within the subgranular zone (SGZ) of the dentate gyrus (DG) generate new neurons that form the granule cell layer during embryonic development and continue to add neurons throughout life. NSCs are multipotent cells, which can undergo self-renewal symmetrically producing daughter cells or differentiate asymmetrically producing astrocytes or intermediate progenitors (Furutachi et al. 2015; Gonçalves et al. 2016). Neurogenesis, maturation, and synaptic integration into the existing network is known to be heavily regulated by cholinergic signaling as shown by selective lesioning and pharmacological studies (Asrican et al. 2016). The targets of acetylcholine are muscarinic and nicotinic acetylcholine receptors found in various cell types throughout the hippocampus (Bertrand et al. 2015). α7 nicotinic acetylcholine receptors (α7 nAChRs) in particular play a critical role in modulation of hippocampal neurogenesis, as the loss of α7 nAChRs results in reduced dendritic length and branch points of newly formed neurons(Campbell et al. 2010). In addition, prior findings in our lab also found that the knockout of α7 nAChRs modulates adult neurogenesis but in a sex-dependent manner (Otto and Yakel 2019). While α7 nAChRs clearly impact neurogenesis the specific cell type mediating these effects is unknown.

GABAergic interneurons play a vital role in distinct stages of DG neurogenesis, regulating NSC quiescence, maturation, and apoptosis (Fernando et al. 2011; Gao et al. 2014; Catavero et al. 2018). α7 nAChRs expressed on GABAergic interneurons are likely to play an important role in mediating these effects, particularly due to their high calcium permeability and impact on second messenger pathways (Haam and Yakel 2017). α7 nAChR-driven signaling has been proposed to directly modulate the level of GABA available to be released via PKA-dependent modulation of GAD67 enzymatic activity (Martin DL 2000; Dajas-Bailador et al. 2002; Wei et al. 2004; Thomsen et al. 2009; Adams et al. 2012; Bates et al. 2014; Cheng and Yakel 2015). In addition, cholinergic signaling is highly responsive to cognitive demand, increasing during encoding of novel information and playing an important role in regulating learning-related synaptic plasticity (Gu and Yakel 2011; Haam and Yakel 2017; Pelkey et al. 2017). α7 nAChRs signaling in interneurons is therefore well positioned to drive learning-induced changes in neurogenesis that occur as a result of exposure to enriched environments and hippocampus-dependent learning (Kempermann et al. 1997; Lemaire et al. 1999; Dalla et al. 2007; Waddell et al. 2011).

The goal of the present study was to assess the structural and functional impact of loss of α7 nAChR expression in GABAergic interneurons utilizing Cre-dependent deletion of the *Chrna*7 gene in a GAD-Cre mouse line (referred to here as *Chrna*7 conditional knockout (cKO) mice). We assessed postnatal hippocampal neurogenesis by crossing the cKO with a GFAP reporter line and conducted a battery of behavioral assays designed to test a broad range of functional domains. Both α7 nAChR-mediated cholinergic signaling and hippocampal neurogenesis have been shown to play important roles in learning and memory processes. Cholinergic signaling supports synaptic plasticity in a timing-dependent manner (Gu and Yakel 2011; Haam and Yakel 2017) and positive allosteric modulators of α7 nAChR have been heavily investigated for their ability to promote cognitive function (Ma and Qian 2019). Ongoing hippocampal neurogenesis plays an important, though hotly debated, role in learning and memory processes (Shors et al. 2002; Nakashiba et al. 2012; Vicidomini et al. 2020). Disruptions of neurogenesis have been shown to impair performance in a wide array of behavioral assays, though deficits are often subtle or require specific task parameters to be observed (Shors et al. 2002; Snyder et al. 2005). We hypothesized that loss α7 nAChRs in GABAergic interneurons would lead to deficits in hippocampal neurogenesis, which could impact behavioral performance. In addition, loss of α7 nAChRs in GABAergic interneurons could impact behavioral performance more directly by eliminating the role this signaling plays in synaptic plasticity and network-level communication. We therefore performed a comprehensive test battery to broadly assess behavioral alterations in *Chrna*7 cKO mice.

## Methods

### Animals

For assessment of postnatal neurogenesis, eight-week-old male and female hGFAP-GFP mice were purchased from Jackson Laboratory (Bar Harbor, ME). These mice were selected as radial glia, astrocytes, and postnatal granule neurons within the DG express GFAP (Garcia et al. 2004). These mice were then crossed with either wildtype (WT; AChRα7_flox_) controls engineered with a loxP site flanking *Chrnα7* or cKO (Gad/AChRα7_flox_) mice expressing GAD-Cre, to allow selective disruption of *Chrnα7* (Hernandez et al. 2014). Numbers were 7 male and 10 female WT mice, and 8 male and 6 female cKO mice. Mice used for immunostaining were group housed in reverse light/dark housing on a 12 hr cycle with food and water supplied *ad libitum*. All procedures were approved and performed in compliance with the NIEHS/NIH Humane Care and Use of Animals Protocols.

For the behavioral battery, the UNC Mouse Behavioral Phenotyping Laboratory utilized 14 male and 13 female WT controls and 10 male and 10 female cKO mice. Mice were 6-8 weeks in age at the start of behavioral testing (see supplemental methods for timeline). All animal care and procedures were conducted in strict compliance with the animal welfare policies set by the National Institutes of Health and by the University of North Carolina at Chapel Hill (UNC), and were approved by the UNC Institutional Animal Care and Use Committee. The NIEHS Neurobehavioral Core utilized 8 male and 8 female WT mice and 8 male and 8 female cKO mice for additional behavioral tests (trace fear conditioning, spontaneous alternation, and object recognition). Mice were 8-10 weeks in age at the start of behavioral testing. These procedures were conducted in strict compliance with the animal welfare policies set by the National Institutes of Health and approved by NIEHS Animal Care and Use Committee. During behavioral assays, animals were acclimated to handling for a minimum of 1 week prior to testing and acclimated to the experimental room for > 30 min prior to initiating testing.

### Tissue Preparation

Mice were deeply anesthetized using 0.2 ml phenobarbital (FatalPlus), transcardially perfused with ice cold 0.1 M phosphate buffer saline (PBS) at pH 7.4 with 0.1% heparin followed by ice cold 4% paraformaldehyde (PFA). Brains were post-fixed overnight at 4°C in 4% PFA, washed in 0.1 M PBS, and cryoprotected in 30% sucrose in 0.1 M PBS at 4°C. Brains frozen in Tissue Freezing Medium (TFM, Triangle Biomedical Sciences) were cryosectioned into 50 μm free-floating coronal sections using a Leica CM305S cryostat. Every sixth section of a hippocampus was processed for immunolabeling. Sections were placed in blocking buffer (0.1 M PBS with 5% goat serum and 0.1% triton x-100) for 2 hours at room temperature or stored at 4°C until immunofluorescence. 50 μm free-floating sections were incubated with primary antibodies overnight at 4° C. Primary antibodies used include guinea pig anti-doublecortin (DCX; 1:500; cat# AB2253, Millipore, Billerica, MA), rabbit anti-ki67 (1:100; cat# 15580 Abcam, Cambridge, MA) chicken anti-GFP (1:10,000; cat# AB13970, Abcam, Cambridge, MA). Negative controls lacking primary were used to confirm specificity of staining. Tissue was triple washed in PBS and incubated in secondary antibody for 2 hours at room temperature or overnight at 4°C. Secondary antibodies were Alexa Fluor 647 goat anti-rabbit (cat# 150087, Abcam, Cambridge, MA), Alexa Fluor 568 goat anti-guinea pig (cat# ab175714, Abcam, Cambridge, MA), Alexa Fluor 488 goat anti-chicken (cat#150169, Abcam, Cambridge, MA). All secondary antibodies were used at 1:500. Tissue was triple washed in PBS with DAPI (2 μg/ml; cat# 508741, Millipore) in the final wash to stain cell nuclei. Tissue was mounted onto SuperFrost Plus slides (Fisher Scientific) using Prolong Diamond Anti-Fade Mounting Media (Molecular Probes).

### Imaging

Tile scan, Z-stack images of 50 μm thick brain sections were collected on a confocal microscope (LSM 710, Carl Zeiss Inc, Oberkochen, Germany) using a 40X objective. For the far-red channel, a 633 nm laser line was used for excitation of an Alexa 647 secondary antibody using a band pass filter of 640-717. For the red channel, a 561 nm dpss laser was used for excitation of Alexa 568 while a 561-639 filter collected the emission. For the green channel, a 488 nm ArKr laser line was used for excitation of the Alexa 488 secondary while a bandpass 503-552 filter was used for collection of the emission signal. For the blue channel, a 405 nm diode laser line was used for excitation of DAPI while a bandpass 415-503 filter was used for the emission. Images were taken with the pinhole of the longest emission wavelength set to 1 airy unit, a zoom setting of 1, and line averaging set to 2.

### Image analysis

All images captured in Zen Black (2012, Carl Zeiss Inc, Oberkochen, Germany) were stitched and imported into an IMARIS (Bitplane, Oxford Industries) analysis arena. 3D images were analyzed using a shape function to limit analysis to the DG within each section. Cells were counted using a Spots function. Where possible, cell counts were automated using the batch feature within IMARIS. Cells counted were confirmed as having a cell nucleus surrounded by the fluorescent marker. Additionally, cells counted were unipolar or bipolar in nature, avoiding astrocytic multipolar morphology. Data was recorded in Excel spreadsheets for further analysis, where densities were calculated. For presentation, images were cropped using IMARIS software, and captured via a snapshot feature. Images were modified solely after analysis by adjusting brightness and contrast to optimize the full dynamic range of the fluorescent signal. For image analysis, raw densities were computed from four or more DG per mouse. Following image analysis by IMARIS and Excel, SPSS (Ill., USA) and/or GraphPad Software (La Jolla California, USA) was used for statistical analysis and graphical illustration. Higher throughput batch cell counting via 3D image analysis software allows for faster gathering of data yet still adds a degree of systematic error. To verify systematic errors inherent in batch counting were not affecting results, cell counts were verified by a separate, blinded count. A student’s t-test was used to analyze two independent samples where opposing hippocampi from a singular mouse provided an internal control. A two-way ANOVA (genotype by sex) was used to analyze main effects and any interactions present. *Post-hoc* analysis included Šídák’s multiple comparison to compare means within relevant rows. Data are presented at mean ± SEM. In all cases a value of p < 0.05 was considered statistically significant.

### Social approach in a 3-chamber choice task

The procedure consisted of three 10-min phases: a habituation period, a test for sociability, and a test for social novelty preference. For the sociability assay, mice were given a choice between being in the proximity of an unfamiliar conspecific (“stranger 1”), versus a non-social novel object (an empty cage). In the social novelty phase, mice were given a choice between the already-investigated stranger 1, versus a new unfamiliar mouse (“stranger 2”). The social testing apparatus was a rectangular, 3-chambered box fabricated from clear Plexiglas. Dividing walls had doorways allowing access into each chamber. An automated image tracking system (Noldus Ethovision) provided measures of time spent in 5-cm proximity to each cage and numbers of entries into each side of the social test box.

At the start of the test, the mouse was placed in the middle chamber and allowed to explore for 10 min, with the doorways into the two side chambers open. After the habituation period, the test mouse was enclosed in the center compartment of the social test box, and an unfamiliar male C57BL/6J adult (stranger 1) was placed in one of the side chambers. The stranger mouse was enclosed in a small Plexiglas cage drilled with holes. An identical empty Plexiglas cage was placed in the opposite side of the chamber. Following placement of the stranger and the empty cage, the doors were re-opened, and the subject was allowed to explore the entire social test box for a 10-min session. At the end of the sociability phase, stranger 2 was placed in the empty Plexiglas container, and the test mouse was given an additional 10 min to explore the social test box.

### Morris water maze

The water maze was used to assess spatial and reversal learning, swimming ability, and vision. The water maze consisted of a large circular pool (diameter = 122 cm) partially filled with water (45 cm deep, 24-26o C), located in a room with numerous visual cues. The procedure involved three different phases: a visible platform test, acquisition in the hidden platform task, and a test for reversal learning (an index of cognitive flexibility).

### Visible platform test

Each mouse was given 4 trials per day, across 2 days, to swim to an escape platform cued by a patterned cylinder extending above the surface of the water. For each trial, the mouse was placed in the pool at 1 of 4 possible locations (randomly ordered), and then given 60 sec to find the visible platform. If the mouse found the platform, the trial ended, and the animal was allowed to remain 10 sec on the platform before the next trial began. If the platform was not found, the mouse was placed on the platform for 10 sec, and then given the next trial. Measures were taken of latency to find the platform and swimming speed via an automated tracking system (Ethovision 15, Noldus, Wageningen, NL).

### Acquisition of spatial learning in the water maze via hidden platform task

Following the visible platform task, mice were tested for their ability to find a submerged, hidden escape platform (diameter = 12 cm). Each animal was given 4 trials per day, with 1 min per trial, to swim to the hidden platform. Criterion for learning was an average group latency of 15 sec or less to locate the platform. Mice were tested until the group reached criterion at 3 days of testing. When criterion was reached, mice were given a one-min probe trial in the pool with the platform removed. Selective quadrant search was evaluated by measuring number of crossings over the platform location, versus the corresponding location in the opposite quadrant.

### Reversal learning in the water maze

Following the acquisition phase, mice were tested for reversal learning, using the same procedure as described above. In this phase, the hidden platform was re-located to the opposite quadrant in the pool. As before, measures were taken of latency to find the platform. On the fourth day of testing, when the criterion for learning was met, the platform was removed from the pool, and the group was given a probe trial to evaluate reversal learning.

### Object recognition assays

Mice were first habituated to the experimental arena (45 cm by 45 cm, 2025 cm2) during Days 1 and 2 for 10 min over two daily sessions. The arena was placed inside a sound attenuating cubicle with an overhead camera to monitor behavior. The light was kept at 5-10 lux and four unique patterned walls were placed just outside the clear walls of the arena to provide distal visual information. On Day 3 the animals were presented with two identical objects (see supplemental methods) placed in opposite corners of the arena for 10 min. During the test phase 24 hours later, one object was moved to the opposite corner, with the side counterbalanced across animals. 24 hours later a novel object recognition assay was performed: the mice were placed back in the arena with two identical novel objects placed equidistant from the walls (6 cm) and from the center of the arena for 10 min. 10 min later one of the objects was replaced with a novel object and the mice were returned to the arena for 10 min (see supplemental methods). The novel versus familiar object was counterbalanced across mice. The amount of time spent exploring the objects was assessed using an automated tracking system (Ethovision) based on parameters that were optimized to closely correlate with human scoring.

### Fear conditioning

Animals were held in an anteroom separated from the testing room to ensure that the animals did not hear testing of other animals for at least 30 min prior to training/testing. Training took place in four identical sound attenuating chambers (Context A; 28 x 21x 21 cm; Med-Associates Inc.). The floor of each chamber consisted of a stainless-steel shock grid (1/2 inch apart) wired to a shock generator and scrambler (Med-Associates Inc.) to deliver foot shocks. On the training day (Day 1) mice underwent trace fear conditioning as follows: mice were placed in Context A (mouse standard grid floor, fan in box on, cleaning solution 75% isopropyl alcohol 3 sprays per box, scent in pan 50% Simple Green 5 sprays) and left to explore for 3 min prior to the first tone presentation (20 s, 75 dB, 2800 Hz). A 2 sec 0.5 mA shock was presented 20 sec later. This sequence was presented 4 more times for a total of 5 tone-shock pairings. After the last shock, animals were left for 2 min and then returned to their home cages. On Day 2 mice were returned to the condition chamber with identical configuration as during training for an 8-min context test. For the tone test on Day 3 the animals were placed in four novel and structurally distinct chambers (Context B-plastic white floor on top of standard grid, Black plastic A-frame, fan off, cleaning solution 75% ethyl alcohol 3 sprays per box, scent in pan Windex 5 sprays). The tone was delivered in the same way as in Context A during training, but with shock omitted.

### Behavioral analysis

Measures were taken by an observer blind to mouse genotype. Behavioral data were first analyzed using two-way or repeated measures ANOVA with genotype and sex as factors. Fisher’s protected least-significant difference (PLSD) tests were used for comparing group means only when a significant F value for genotype was determined in the ANOVA. Within-genotype comparisons were used to determine side preference in the three-chamber test for social approach and quadrant selectivity in the Morris water maze. For all comparisons, significance was set at p < 0.05.

## Results

### Assessment of postnatal hippocampal neurogenesis

We found that the selective knockout of *Chrna7* (cKO mice) within GABAergic interneurons resulted in a sex by genotype interaction in postnatal neuron density within the DG as marked by GFAP and DAPI positive cells with neuron-like morphology (F_1,27_ = 12.9, p = 0.013; **Fig. 1 *A-P***). Postnatal neuron density was reduced in male cKO mice by 42 ± 10% compared to male WT mice, but there no differences were detected between female genotypes. Within WT mice, we observed fewer postnatal neurons in females relative to males, but no sex difference was observed in the cKO mice. There was no significant difference in the numbers of immature neurons or in actively dividing cells in either male or female mice (ki67: F_1,27_ = 1.762, p = 0.196, **Fig. 1 *R***; DCX: F_1,27_ = 3.919, p = 0.058, **Fig 1 *S***).

**Fig. 1.**
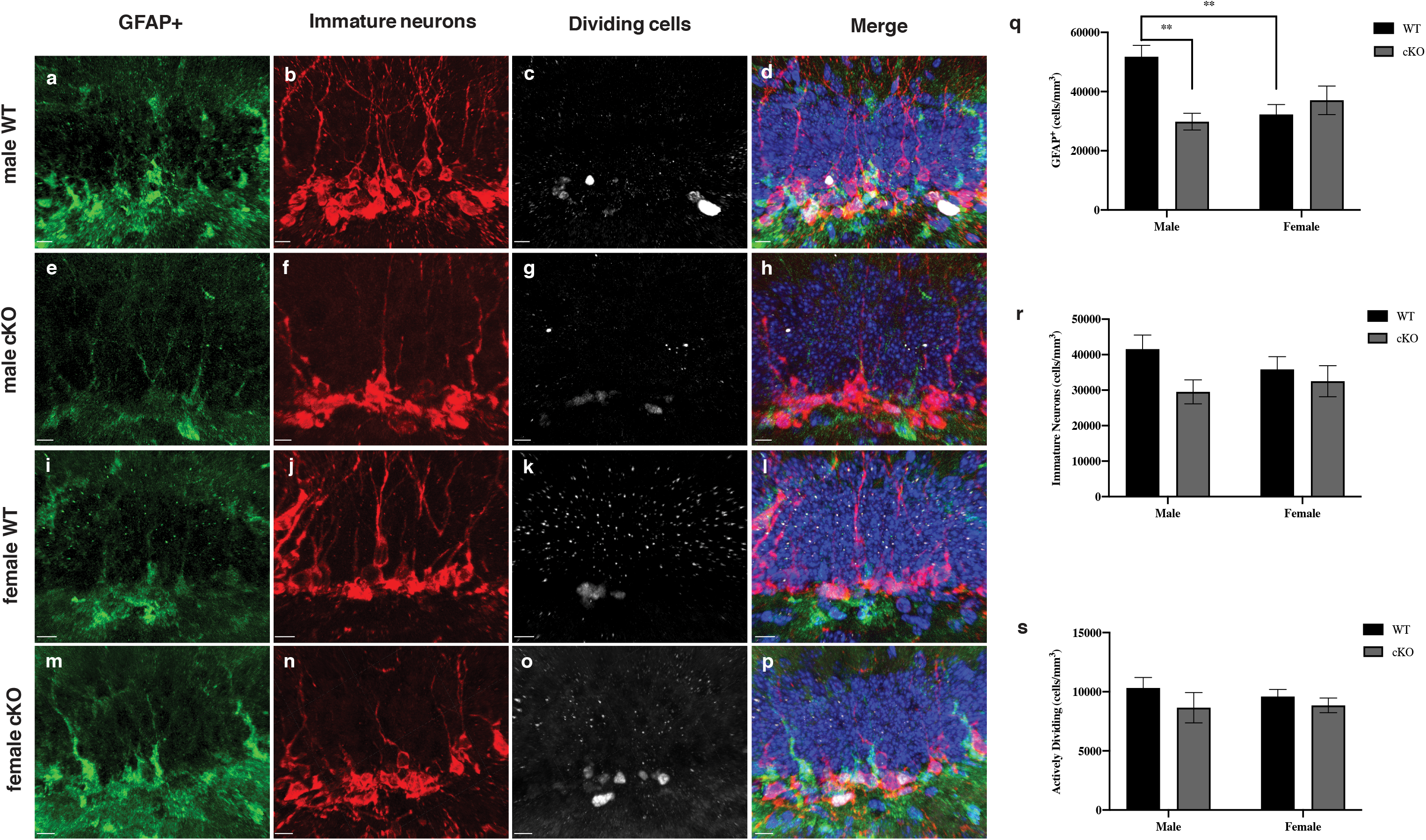
Loss of α7 nAChRs in GABAergic interneurons reduces postnatal granule cells in male mice. ***a-p*** 40x 3D images of the DG from hGFAP-GFP x α7 nAChR_flox_ x GADCre mice to visualize radial glia, astrocytes, and postnatal granule neurons (***a-d*** male WT; ***e-h*** male cKO; ***i-l*** female WT; ***m-p*** female cKO). Expression of GFP was amplified using ck-antiGFP (1:10,000). Mice were also probed for immature neuron expression using gp-antiDCX (1:500) and actively dividing cells using rb-antiki67 (1:400). Cell nuclei were marked using DAPI (1:500). Scale bar = 10 μm. ***q-s*** GFAP+, DCX+ and Ki67 + cell densities were measured, and plotted against sex and genotype. **p < 0.01. Data expressed as average cell count ± SEM

### Overall behavioral findings

When examining impacts of postnatal neurogenesis at the phenotypic level, several behavioral assays were performed. The cKO mice had increased body mass in comparison to the WT mice at most time points during the behavioral study (**Supplemental Fig. 1**). No differences were found in the following tasks: Marble burying, open arm entries in elevated plus maze and olfactory test (**Supplemental Table 1**), rotarod (**Supplemental Fig. 2**), startle responses following acoustic stimuli (**Supplemental Fig. 3)**, prepulse inhibition of acoustic startle response (**Supplemental Fig. 4**), open field distance and rearing within male WTs and cKOs (**Supplemental Fig. 6**), and spontaneous alternation (**Supplemental Fig. 8**). Overall, the *Chrnα7* mutation did not lead to overt changes in health or motor ability for all tasks (**Supplemental Fig. 10**).

#### Three chamber social approach test

An ANOVA on time in proximity to each cage indicated a significant main effect of sex (F_1,40_ = 19.53, p < 0.0001); therefore, data from males and females were separately analyzed. As shown in **Fig. 2 A,B**, both WT and cKO mice had robust preference for spending more time in proximity to the stranger mouse, versus the empty cage (within-genotype comparisons following repeated measures ANOVA, significant effect of side in males, F_1,19_ = 98.21, p < 0.0001; and females, F_1,21_ = 65.64, p < 0.0001). However, in the male groups, a different pattern was observed during the test for social novelty (**Fig. 2 C**). In contrast to WT, the male cKO mice failed to exhibit significant preference for the newly-introduced stranger mouse (main effect of genotype, F_1,19_ = 12.79, p = 0.002; genotype by side interaction, F_1,19_ = 6.32, p = 0.021). Data were removed for one outlier female cKO mouse that remained in proximity to the second stranger mouse for 466 sec. No significant effects of genotype were observed in the female groups during the social novelty test (**Fig. 2 D**). As shown in **Supplemental Fig. 5**, the WT and cKO groups had similar numbers of entries during the social approach test, indicating the significant differences in the male groups were not due to altered activity or exploration in the cKO mice.

**Fig. 2.**
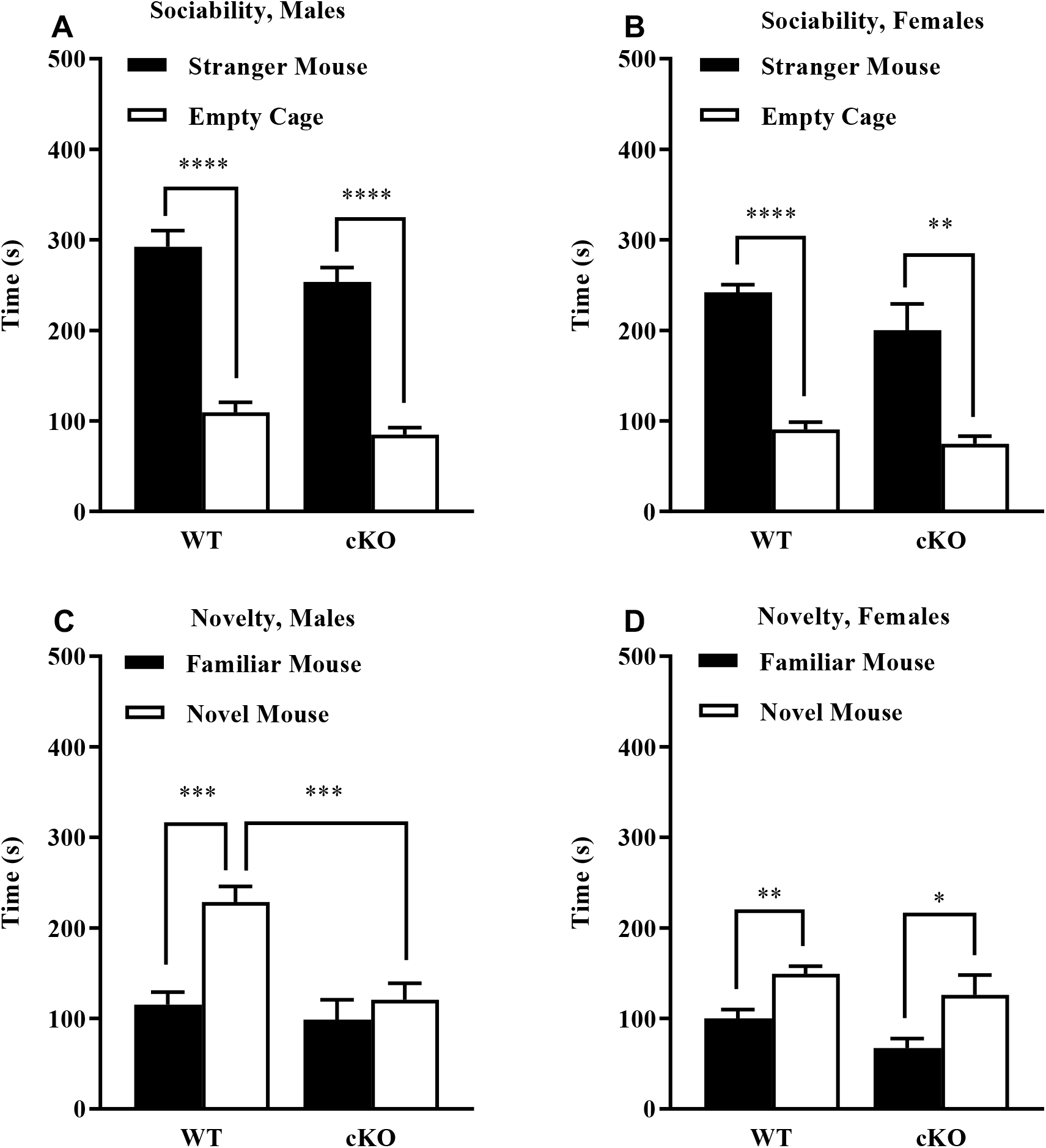
Reduced social novelty preference in cKO male mice during a 3-chamber choice task. Measures were taken of time spent in proximity to each cage during the test for sociability ***a,b*** and preference for social novelty ***c,d***. Data were lost from one male CKO mouse, due to experimenter error. Data were removed for one outlier female cKO mouse that remained in proximity to novel mouse for 466 sec. **p < 0.01, ***p < 0.001, ****p < 0.0001. Data expressed as mean ± SEM

#### Morris water maze

The WT and cKO mice showed high proficiency in the visible platform phase of the water maze test (**Supplemental Table 2**). All experimental groups had mean latencies of less than 10 sec by the second day of visible platform testing. However, a significant main effect of genotype was found for swimming speed on the first day of testing for each phase (F_1,40_ = 15.04, p = 0.0004; no effect of sex). Further analysis indicated the female cKO mice had lower swimming speeds than the female WT at each time point (**Supplemental Table 2**). The WT and cKO mice demonstrated similar rates of learning, measured by latency to reach the escape platform, during acquisition of the hidden platform task (**Fig. 3 A,B**), and the reversal phase (**Fig. 3 C,D**).

**Fig. 3.**
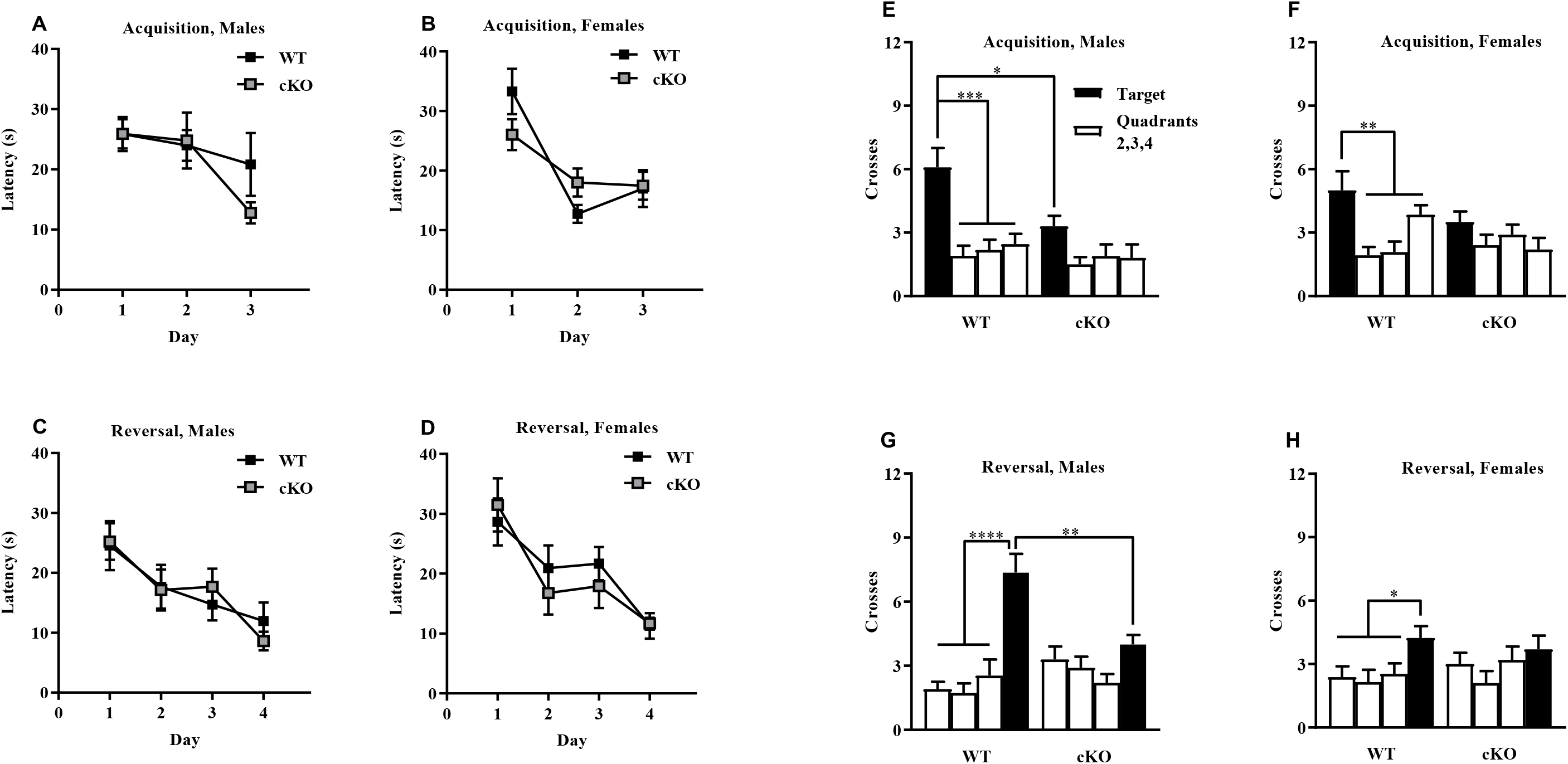
Normal acquisition learning in a hidden platform task but lack of quadrant preference in cKO mice during probe trials. Criterion for learning was a group average of 15 sec to find the escape platform ***a-d***. Mice were given a one-min probe trial without the platform following an initial 3-day acquisition of the hidden platform task ***e,f*** and a subsequent 4-day reversal phase ***g,h***. ‘Target’ indicates the quadrant where the platform was located in each phase. *p < 0.05, **p < 0.01, ***p < 0.001, ****p < 0.0001. Data expressed as mean ± SEM of four trials per day in the Morris water maze

#### Probe trials in water maze test

At the end of the acquisition and reversal phases, mice were given one-min probe trials with the platform removed. An ANOVA on number of platform-location crossings during the acquisition probe trial revealed significant effects of genotype and interaction (main effect, F_1,40_ = 8.16, p = 0.007; genotype by quadrant interaction, F_3,120_ = 3.55, p = 0.017; no effect of sex). As shown in **Fig. 3 E** and **F**, male and female WT mice demonstrated significant quadrant selectivity measured by number of crossings. The cKO mice failed to exhibit a higher number of crossings across the target location. In male groups, the cKO mice had significantly fewer crossings than WT across the target (*post-hoc* test following main effect of genotype in male groups, p = 0.007). A similar pattern was observed in the probe test following the reversal phase (**Fig. 3 G** and **H**), wherein the cKO mice again failed to show quadrant selectivity (genotype by quadrant interaction, F_3,120_ = 4.71, p = 0.004; no effect of sex). In the male groups, the cKO mice had decreased numbers of crossings over the new location for the platform (*post-hoc* test following genotype by quadrant interaction in male groups, p = 0.001).

#### Object Tasks

On Day 3 during object familiarization, cKO mice spent significantly less total time investigating the novel objects (F_1,32_ = 8.097, p = 0.008; **Fig. 4 A**). Despite this overall reduction in investigation, the cKO mice demonstrated a similar preference for the displaced object when calculated as percent investigation (displaced/total investigation versus stationary/total investigation), which normalizes for the total amount of investigation time (**Fig. 4 B**). Overall effect of object (F_1,25_ = 13.87, p = 0.001), within cKO mice: p = 0.015 and within WT mice: p = 0.029. During the novel object recognition task. we again saw an overall reduction in object investigation in cKO mice (F_1, 24_ = 4.455, p = 0.045; **Fig. 4 C** and **D**) and a similar preference for the novel object as measured by percent investigation (effect of object in WT: p = 0.022; effect of object in cKO: p = 0.005). See **Supplemental Fig. 7** for remaining SOR and NOR findings and **Supplemental Fig. 9** for objects.

**Fig. 4.**
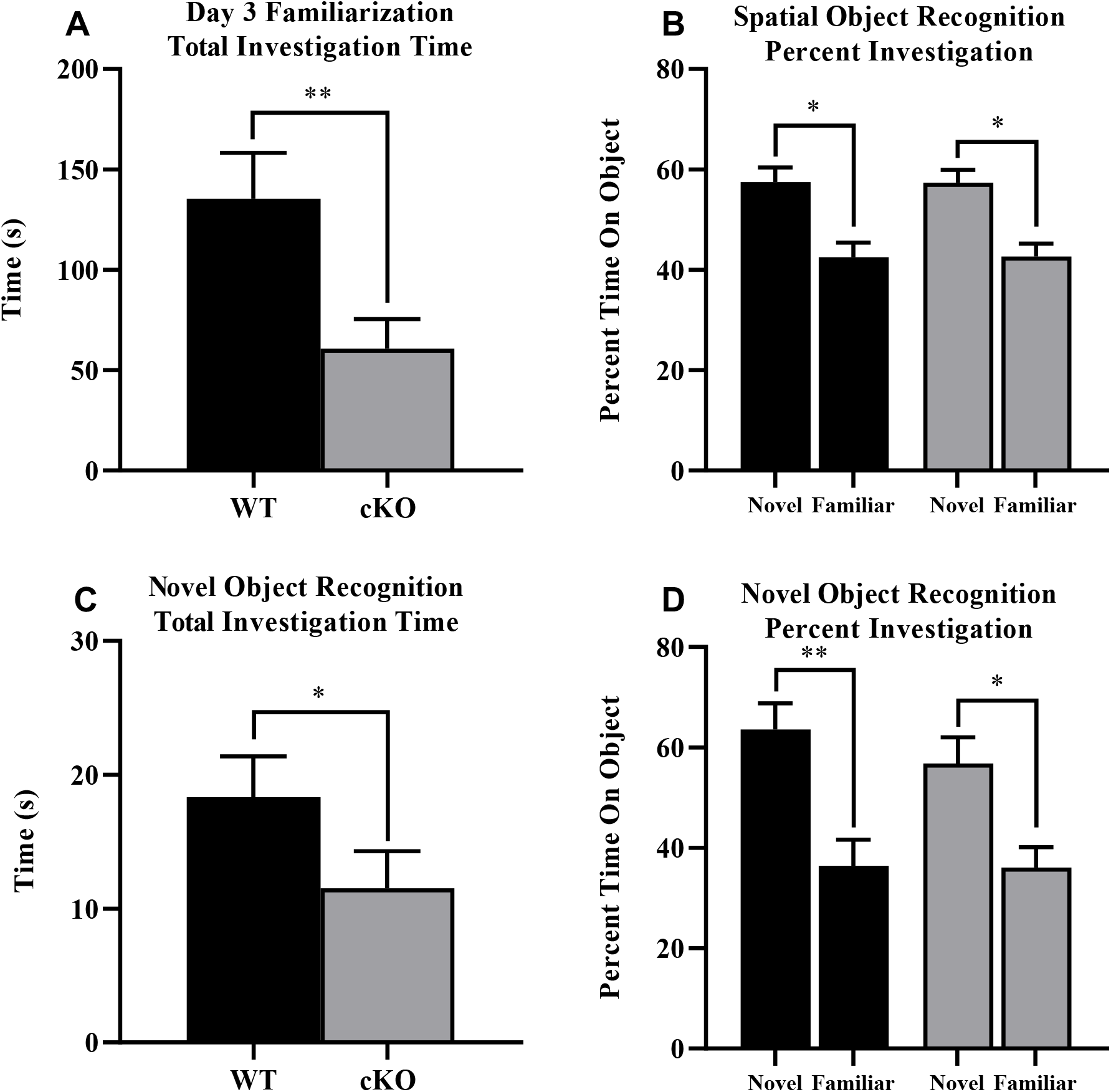
Decreases in object investigation during SOR and NOR. Measures were taken of total object investigation familiarization time during SOR as an average of time and percentage ***a,b***. Total investigation time during NOR was recorded and plotted as an average of time and percentage ***c, d***. *p < 0.05, **p < 0.01, ***p < 0.001, ****p < 0.0001. Data expressed as mean ± SEM

#### Fear Conditioning

Freezing increased across the session during initial acquisition training, but did not differ by genotype (effect of time: F_10,280_ = 1.617, p = 0.101; genotype by time interaction: F_10,280_ = 0.299, p = 0.981; Fig. 5 A). No differences in shock reactivity were observed (F_1,28_ = 0.029, p = 0.866). No genotype or sex differences were detected during context testing (F_1,28_ = 0.503, p = 0.484; Fig. 5 B). No difference in baseline (F_1,28_ = 0.148, p = 0.704), tone (F_1,28_ = 0.720, p = 0.403), or post-tone freezing (F_1,28_ = 0.178, p = 0.676) was detected (Fig. 5 C).

**Fig. 5.**
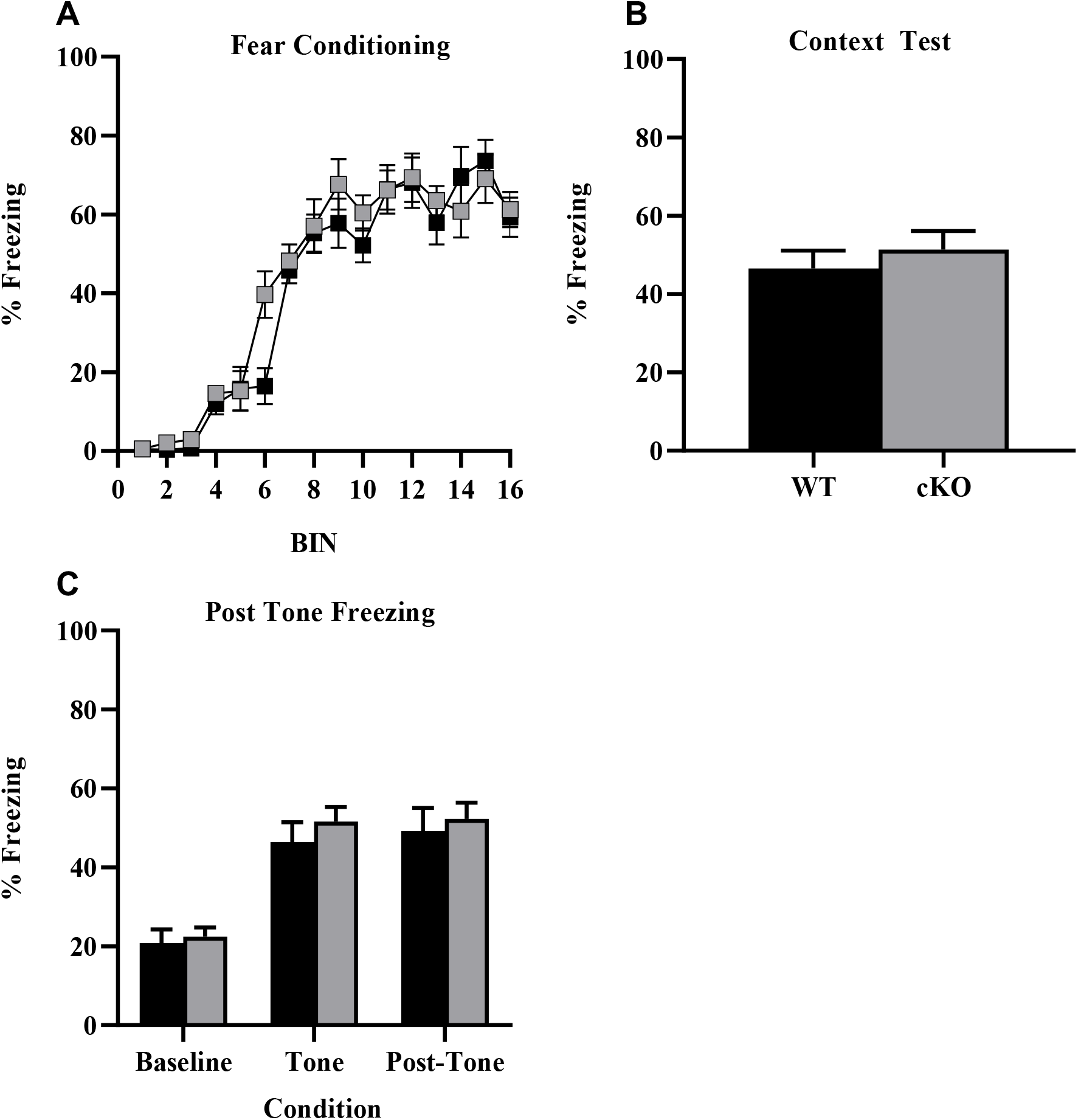
Lack of differences in fear conditioning and context test. Measures were taken of total freezing during sixteen trials across training ***a***. Total percentage freezing during context test was recorded ***b*** with baseline, tone, and post-tone freezing ***c***. Data expressed as average percentage freezing ± SEM

## Discussion

Our findings indicate that α7 nAChRs expressed in GABAergic interneurons play an important, sex-dependent role in the regulation of postnatal hippocampal neurogenesis and behavior. Male α7 cKO mice showed decreased postnatal neurogenesis, as measured by GFAP+ cell density in the DG, whereas female α7 cKO were unchanged relative to female WT mice. Proliferation, as measured by Ki67+ density, did not change, and there was no significant difference in immature neurons, as measured by DCX+ cell density. Additionally, within WT mice we observed fewer GFAP+ neurons in female mice relative to males, suggesting a major underlying sex difference in the level of postnatal neurogenesis, consistent with some prior findings (Juraska et al. 1985; Falconer and Galea 2003; Brydges et al. 2018; Yagi and Galea 2019). In a battery of behavioral tests, the most notable deficits were a male-specific deficiency in social recognition and impaired spatial learning that was strongest in the male cKO mice.

The behavioral changes in α7 cKO mice were highly selective indicating specific functional roles for *Chrna7* expressed in interneurons. Disruption of *Chrna7* did not lead to changes in anxiety-like behavior in the elevated plus maze or marble burying tests. In the female groups, the cKO mice showed reduced rearing in a 1-hr open field test, and slower swimming speeds in the water maze, in comparison to WT females. Although these changes suggest a possible motor impairment, both male and female cKO mice were proficient in a rotarod test for motor coordination and in the visible platform test in the water maze. The selective deletion of *Chrnα7* did not lead to alterations in prepulse inhibition of acoustic startle responses, indicating the cKO mice did not model the deficits in sensorimotor gating associated with schizophrenia. Increased sensitivity to thermal stimuli was present in both male and female cKOs, consistent with previously reported findings of α7 nAChRs mediating nociception by reducing pain evoked by heat (Rowley et al. 2008). The augmentation in weight for cKOs warrants further study as α7 nAChRs have been shown to be important in metabolic function, including cancer progression and oxidative stress (Bertrand et al. 2015). Oxidative stress accumulation in adipose tissue has been associated with insulin resistance, a biomarker for weight gain or obesity (Aroor and DeMarco 2014), which could potentially involve α7 nAChR signaling.

In a 3-chamber test for sociability, both WT and cKO mice had similar preference for proximity to a stranger mouse, versus a non-social novel object. However, in a test for social novelty preference, the male cKO mice failed to demonstrate the typical shift in attention towards a newly-introduced stranger mouse, indicating a specific deficit in social recognition. This male-specific behavioral difference mirrors the male-specific deficit in postnatal neurogenesis, which could indicate a link between the two. Decreases in social approach have been correlated with decreases in DG volume, reported in both humans and in mouse models (Kalman and Keay 2017), and deletion of the α4 GABA receptor subunit has been shown to impair social recognition (Fan et al. 2020). Furthermore, studies have shown male-specific social and behavioral deficits following exposure to the pesticide chlorpyrifos (Lan et al. 2019), which is known to target the α7 nAChR (Slotkin et al. 2004). Given the greater preponderance of autism in males, these findings suggest GABAergic and cholinergic signaling may contribute to social deficits by impacting postnatal neurogenesis in a sex-dependent manner. Evident sex differences with regards to social housing and chronic stress have been correlated to levels of neurogenesis, where male rats experience a reduction and females experience an increase in proliferating cell counts within granule cell layer (Westenbroek et al. 2004).

In the Morris water maze task, α7 cKO mice showed impaired preference for the platform location during the probe trial indicating a deficit in spatial learning and memory. This deficit is strongest in the male cKO, although female cKO also failed to show a spatial bias for the target quadrant. Female controls showed a less pronounced spatial bias relative to control males, consistent with some prior evidence of relative spatial deficits in females (Yagi and Galea 2019). This reduced bias likely created a floor effect such that the target quadrant preference in the female cKO mice was not significantly reduced relative to female controls. The overall sex-dependent pattern in spatial performance actually mirrors the differences that we observed in the level of post-natal neurogenesis: a relative reduction in control females relative to control males and a specific reduction in male cKO relative to male controls. These findings mirror our prior results which showed full body α7 KO produced sex dependent alterations reduction in hippocampal neurogenesis and a male-specific deficit in spatial pattern separation (Otto and Yakel 2019). Together these findings suggest a possible relationship between the level of postnatal neurogenesis and overall spatial performance. Postnatal hippocampal neurogenesis has been shown to have little role in short term spatial memory in the water maze (Snyder et al. 2005), however it does impact spatial search strategies (Yu et al. 2019). The spatial impairment in the cKO mice could also be due to more direct effects of aberrant cholinergic signaling on encoding, retrieval and task performance. Prior studies have shown that hilar interneurons play a critical role in water maze performance, which argues that impaired cholinergic modulation of interneurons could mediate the observed spatial deficit in α7 cKO mice (Andrews-Zwilling et al. 2012).

In contrast to the spatial learning deficit, α7 cKO mice showed normal hippocampus-dependent performance in a number of other behavioral assays indicating that α7-dependent cholinergic modulation of interneurons plays a very specific role in hippocampus-dependent learning and memory. Spontaneous alternation, considered a measure of spatial working memory, was normal, indicating the deficit may be more specific to long term retention of the spatial location of the escape platform. In contrast to this, however, object location performance was intact, suggesting that some forms of long-term spatial learning and memory are unaffected in the α7 cKO mice. This task was complicated however, by a decrease in the overall time the α7 cKO mice spent investigating the objects. Despite this reduction, they showed a similar preference for the displaced object when normalizing by the total investigation time. The exact mechanisms underlying this reduced object investigation are unclear, however prior studies have linked general reactivity to novelty to hippocampal neurogenesis (Lemaire et al. 1999) and to differences in investigation of novel objects, in particular (Denny et al. 2012).

No differences were found in trace or contextual fear conditioning, again confirming, the specificity of the spatial deficits. This is surprising given the well-established role for cholinergic and GABAergic signaling in hippocampus-dependent forms of fear conditioning (Moore et al. 2010; Cushman et al. 2014; Hersman et al. 2017). The increased thermal sensitivity suggests that pain sensitivity to the shock may have been altered in the α7 cKO mice, however the magnitude of the activity burst response to the shock did not differ. One possibility for the lack of an effect is that nicotinic manipulations have been shown to have opposing roles in the dorsal vs. ventral hippocampus (Kenney et al. 2012). The effects of losing α7-mediated cholinergic modulation of GABAergic interneurons in the whole hippocampus might therefore be counteracted by differential dorsal versus ventral effects.

Overall, we have shown that α7 nAChRs on GABAergic interneurons play a sex dependent role in regulating postnatal neurogenesis, social recognition, and spatial learning and memory. Loss of homomeric α7 nAChRs in GABAergic interneurons appears to impact very specific domains of hippocampal function as other hippocampus-dependent measure were unaffected. The finding of a male-specific role for α7 nAChRs in the modulation of both postnatal neurogenesis and social behavior may provide a potential mechanism for why males are more sensitive to developmental neurotoxicants that target the cholinergic system. Although further research is required, these findings argue that greater insight into this potential mechanism could lead to novel treatments for neuropsychiatric and neurodegenerative disorders.

## Acknowledgements

We would like to acknowledge Patricia Lamb for creation and maintenance of mouse lines, and Charles J. Tucker for assistance with confocal microscopy. This research was supported by the Intramural Research Program of the NIH, National Institute of Environmental Health Science.

