## Supplemental Figures & Tables for "Loss of α7 nicotinic acetylcholine receptors in GABAergic interneurons causes sex-dependent impairments in postnatal neurogenesis and cognitive and social behavior"

### Supplemental Information

### Methods

#### Elevated plus maze test.

This test was used to assess anxiety–like behavior, based on a natural tendency of mice to actively explore a new environment, versus a fear of being in an open area. Mice were given one five-min trial on the plus maze, which had two walled arms (the closed arms, 20 cm in height) and two open arms. The maze was elevated 50 cm from the floor, and the arms were 30 cm long. Mice were placed on the center section (8 cm x 8 cm) and allowed to freely explore the maze. Measures were taken of time on, and number of entries into, the open and closed arms.

#### Open field test.

Exploratory activity in a novel environment was assessed by a one-hour trial in an open field chamber (41 cm x 41 cm x 30 cm) crossed by a grid of photobeams (VersaMax system, AccuScan Instruments). Counts were taken of the number of photobeams broken during the trial in five-min intervals, with measures taken of locomotion (total distance traveled), rearing movements, and time spent in the center region of the open field, an index of anxiety-like behavior.

#### Marble-burying assay.

Mice were tested in a Plexiglas cage located in a sound-attenuating chamber with ceiling light and fan. The cage contained 5 cm of corncob bedding, with 20 black glass marbles (14 mm diameter) arranged in an equidistant 5 X 4 grid on top of the bedding. Subjects were given access to the marbles for 30 min. Measures were taken of the number of buried marbles (two thirds of the marble covered by the bedding).

#### Rotarod test.

Subjects were tested for motor coordination and learning on an accelerating rotarod (Ugo Basile, Stoelting Co., Wood Dale, IL). For the first test session, mice were given three trials, with 45 sec between each trial. Two additional trials were given 48 hours later. Rpm (revolutions per min) was set at an initial value of 3, with a progressive increase to a maximum of 30 rpm across 5 min (the maximum trial length). Measures were taken for latency to fall from the top of the rotating barrel.

#### Acoustic startle test.

This procedure was used to assess auditory function, reactivity to environmental stimuli, and sensorimotor gating. The test was based on the reflexive whole-body flinch, or startle response, that follows exposure to a sudden noise. Measures were taken of startle magnitude and prepulse inhibition, which occurs when a weak pre-stimulus leads to a reduced startle in response to a subsequent louder noise. Subjects were given 2 tests, the first at 11-13 weeks of age, and the sec test at 17-19 weeks of age.

For each test, mice were placed into individual small Plexiglas cylinders within larger, sound-attenuating chambers. Each cylinder was seated upon a piezoelectric transducer, which allowed vibrations to be quantified and displayed on a computer (San Diego Instruments SR-

Lab system). The chambers included a ceiling light, fan, and a loudspeaker for the acoustic stimuli. Background sound levels (70 dB) and calibration of the acoustic stimuli were confirmed with a digital sound level meter (San Diego Instruments). Each session consisted of 42 trials, that began with a five-min habituation period. There were 7 different types of trials: the no-stimulus trials, trials with the acoustic startle stimulus (40 msec; 120 dB) alone, and trials in which a prepulse stimulus (20 msec; either 74, 78, 82, 86, or 90 dB) occurred 100 ms before the onset of the startle stimulus. Measures were taken of the startle amplitude for each trial across a 65-msec sampling window, and an overall analysis was performed for each subject's data for levels of prepulse inhibition at each prepulse sound level (calculated as 100 - ((response amplitude for prepulse stimulus and startle stimulus together / response amplitude for startle stimulus alone) x 100).

**Buried food test.**

Several days before the olfactory test, an unfamiliar food (Froot Loops, Kellogg Co., Battle Creek, MI) was placed overnight in the home cages of the mice. Observations of consumption were taken to ensure that the novel food was palatable. Sixteen to twenty hours before the test, all food was removed from the home cage. On the day of the test, each mouse was placed in a large, clean tub cage (46 cm L x 23.5 cm W x 20 cm H), containing paper chip bedding (3 cm deep), and allowed to explore for five min. The animal was removed from the cage, and one Froot Loop was buried in the cage bedding. The animal was then returned to the cage and given fifteen min to locate the buried food. Measures were taken of latency to find the food reward.

**Wire-hang test.**

Each mouse was placed on a large metal cage lid. The lid was gently shaken to induce the mouse to grip the metal grid. The cage top was then inverted over a large foam pad, and latency for the mouse to fall from the lid was recorded. The maximum trial length was 60 sec.

**Hotplate test.**

Individual mice were placed in a tall plastic cylinder located on a hotplate, with a surface heated to 55$^{o}$ C (IITC Life Science, Inc., Woodland Hills, CA). Reactions to the heated surface, including hind paw lick, vocalization, or jumping, led to immediate removal from the hotplate. Measures were taken of latency to respond with a maximum test length of 30 sec.

**Spontaneous alternation.**

Mice were placed in a novel 3-arm maze and allowed to explore for 8 min. The arms measured 38.5 cm long, 3 cm wide, 13 cm high, and were oriented at 60° angles from each other. The maze was placed in the center of black curtained area with a camera mounted over the center of the maze. Large white cues on the curtains provided distal visual information. Lighting was held at 5 lux. Mice were placed at the end of the start arm of the Y-maze within a curtained area. The sequence of arm entries and total distance moved were recorded. Alternations were counted as making a visit to each of the three arms in succession. Percent alternation was calculated as the number of alternations divided by the total possible.

### Results

**Elevated plus maze.** As shown in **Supplemental Table 1**, the WT and cKO groups had similar percent time on the open arms of the maze, and similar percent entries, suggesting the loss of *Chrnα7* did not change anxiety-like behavior. However, a significant main effect of genotype was found for total number of arm entries during the test, a measure of general activity ($F_{(1,43)}$=12.87, p=0.0008; no effect of sex]. Separate ANOVAs confirmed that both male and female cKO mice made significantly more entries than WT controls on the elevated plus maze.

**Sensitivity to thermal stimulus.** As shown in **Supplemental Table 1**, both the male and female cKO mice had increased sensitivity to the thermal stimulus (*post-hoc* tests following main effect of genotype, $F_{1,40}$ = 9.07, p = 0.0045; no effect of sex).

**Marble-burying assay.** Both WT and cKO mice buried comparable numbers of marbles during the test (**Supplemental Table 1**).

**Wire-hang test** for grip strength, all of the mice except one were able to remain suspended from a cage lid for the maximum trial time of 60 sec.

**Health Status.** The cKO mice had increased body mass in comparison to the WT mice at most time points during the behavioral study (**Supplemental Fig. 1**). These significant differences were observed in the male groups (main effect of genotype, $F_{1,19}$ = 6.36, p = 0.021; genotype by time interaction, $F_{2,38}$ =14.49, p < 0.0001) and the female groups (main effect of genotype, $F_{1,21}$ = 21.8, p = 0.0001; genotype by time interaction, $F_{2,42}$ = 8.44, p = 0.0008).

**Open field results.** As shown in **Supplemental Fig. 6**, no differences were observed between the male WT and cKO mice for activity in the open field. In the female groups, the cKO mice had decreased locomotion, measured by distance traveled, starting near the middle of the session (main effect of genotype, $F_{1,21}$ = 4.96, p = 0.037). Further, the female cKO group had a striking reduction in rearing responses at all but the first interval during the one-hour test (main effect of genotype, $F_{1,21}$ = 19.62, p = 0.0002; genotype by time interaction, $F_{11,231}$ = 2.57, p = 0.004). In contrast, no genotype effects were found for time spent in the center region (summary data in **Supplemental Table 1**).

**Spontaneous Alternation.** The number of spontaneous alternations between radial arm visits demonstrated a trend towards a reduction in cKO mice (main effect of genotype, $F_{1,28}$ = 3.762, p = 0.062; **Supplemental Fig. 8A**). However, no difference in percent alternation was observed (main effect of genotype, $F_{1,28}$ = 0.049, p = 0.826; **Supplemental Fig. 8B**).

**Supplemental Fig 1. Increased weight in cKO mice.** Measures were taken of total mass per mouse while age was approximate number of weeks ***a,b.*** *p < 0.05, ***p < 0.001, ****p < 0.0001. Data expressed as mean ± SEM

**Supplemental Fig 2. No differences in latency to fall from an accelerating rotarod**. Maximum trial length was 300 sec. Trials 4 and 5 were given 48 hours after the first 3 trials ***a,b***. Data expressed as mean ± SEM

**Supplemental Fig 3. No differences in magnitude of startle responses following presentation of acoustic stimuli.** Trials included no stimulus (No S) trials and acoustic startle stimulus (AS; 120 dB) alone trials. Test 1 was given when mice were 11-13 weeks of age, and Test 2, at 17-19 weeks of age ***a-d***. Data expressed as mean ± SEM

**Supplemental Fig 4. No differences in prepulse inhibition of acoustic startle responses.** Percent inhibition was measured and plotted against startle noise. Test 1 was given when mice were 11-13 weeks of age, and Test 2, at 17-19 weeks of age ***a-d***. Data expressed as average percent inhibition ± SEM

**Supplemental Fig 5. No differences in entries into each side during a 3-chamber choice task.** Sociability procedure consisted of three 10-min phases. Initial exposure to stranger followed by more complex social discrimination ***a-d*** . Data expressed as mean ± SEM for a 10-min test

**Supplemental Fig 6**. **Reduced distance traveled and rearing movements in a novel environment in cKO females.** Distanced traveled and number of rearing movements were recorded as averages and plotted ***a-d***. Data expressed as mean ± SEM for each group for a one-hour test session. *p < 0.05, ***p < 0.001, ****p < 0.0001

**Supplemental Fig 7. Decrease in object investigation during object location.** Measures were taken of distance traveled in first day, along total object investigation familiarization time during SOR as an average of time and percentage ***a,b*** Total investigation time during NOR was recorded and plotted as an average of time and percentage ***c,d*** while broken down by sex ***e,f.*** *p < 0.05, **p < 0.01, ***p < 0.001, ****p < 0.0001. Data expressed as mean ± SEM

**Supplemental Fig 8. No differences in spontaneous alternation** Measurements were taken as number of arm visits with a trend towards reduction in entries and no difference in percent alternation ***a,b***. Data expressed as average number of arm visits ± SEM

**Supplemental Fig 9. NOR and SOR** Objects 1 and 2 were used for novel object recognition with counterbalancing, while 3 was used in spatial object recognition.

**Supplemental Fig 10. Timeline** provided by UNC mouse behavioral phenotyping Laboratory

**Supplemental Table 1.** Performance in tests for exploration and anxiety-like behavior (elevated plus maze, open field, and marble-burying assays), olfactory ability, and sensitivity to a thermal stimulus.

| | Males | | Females | |
|---|---|---|---|---|
| | WT | cKO | WT | cKO |
| Elevated plus maze (5 min) | | | | |
| Percent open arm time | 21 ± 4 | 15 ± 2 | 18 ± 4 | 11 ± 2 |
| Percent open arm entries | 33 ± 2 | 29 ± 2 | 30 ± 3 | 27 ± 2 |
| Total number of entries | 13 ± 2 | 20 ± 2* | 15 ± 1 | 21 ± 3* |
| Open field (1 hr) | | | | |
| Total time in the center region (sec) | 233 ± 26 | 278 ± 50 | 195 ± 19 | 160 ± 42 |
| Marble-burying | | | | |
| Number buried in 30 min. | 17 ± 0.8 | 16 ± 0.6 | 17 ± 0.3 | 16 ± 0.7 |
| Olfactory test (15 min) | | | | |
| Latency to find buried food (sec) | 93 ± 27 | 139 ± 43 | 63 ± 14 | 187 ± 78 |
| Hotplate (30 sec) | | | | |
| Latency to respond (sec) | 29 ± 0.7 | 26 ± 1.4* | 28 ± 0.9 | 24 ± 1.7* |

*$p < 0.05$

**Supplemental Table 2.** Vision and swimming ability in the Morris water maze. Data are means (± SEM) of 4 trials per day.

| | Males | | Females | |
|---|---|---|---|---|
| | WT | cKO | WT | cKO |
| Visible platform, latency to escape (sec) | | | | |
| Day 1 | 25 ± 5 | 17 ± 2 | 20 ± 3 | 22 ± 4 |
| Day 2 | 9 ± 2 | 9 ± 1 | 8 ± 1 | 9 ± 2 |
| Swim speed (cm/sec) | | | | |
| Day 1, visible platform test | 17 ± 0.6 | 17 ± 0.9 | 18 ± 0.7 | 15 ± 0.7* |
| Day 1, acquisition | 20 ± 0.6 | 18 ± 0.7 | 20 ± 0.7 | 18 ± 0.5* |
| Day 1, reversal | 19 ± 0.6 | 18 ± 0.6 | 21 ± 0.9 | 18 ± 0.7* |

*$p < 0.05$

**Supplemental Fig. 1**

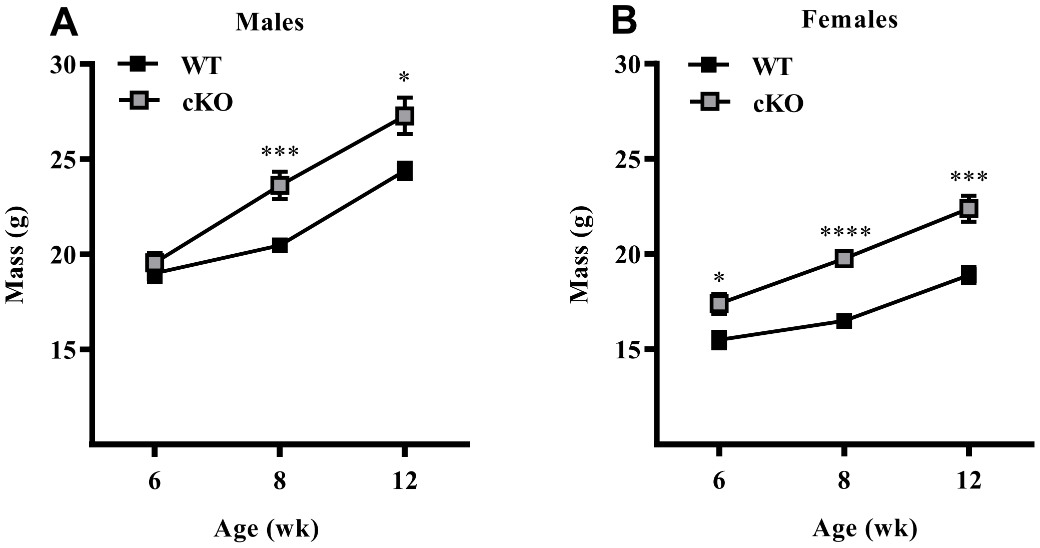

**Supplemental Fig. 2**

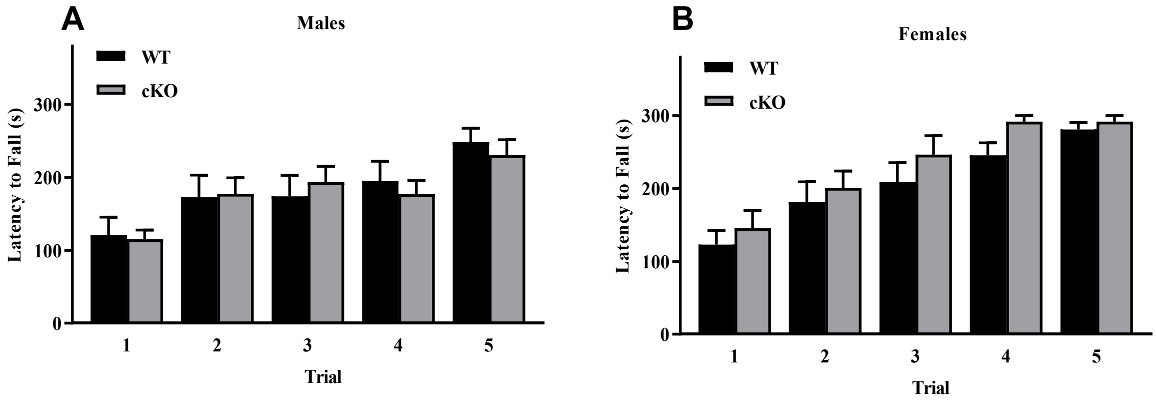

**Supplemental Fig. 3**

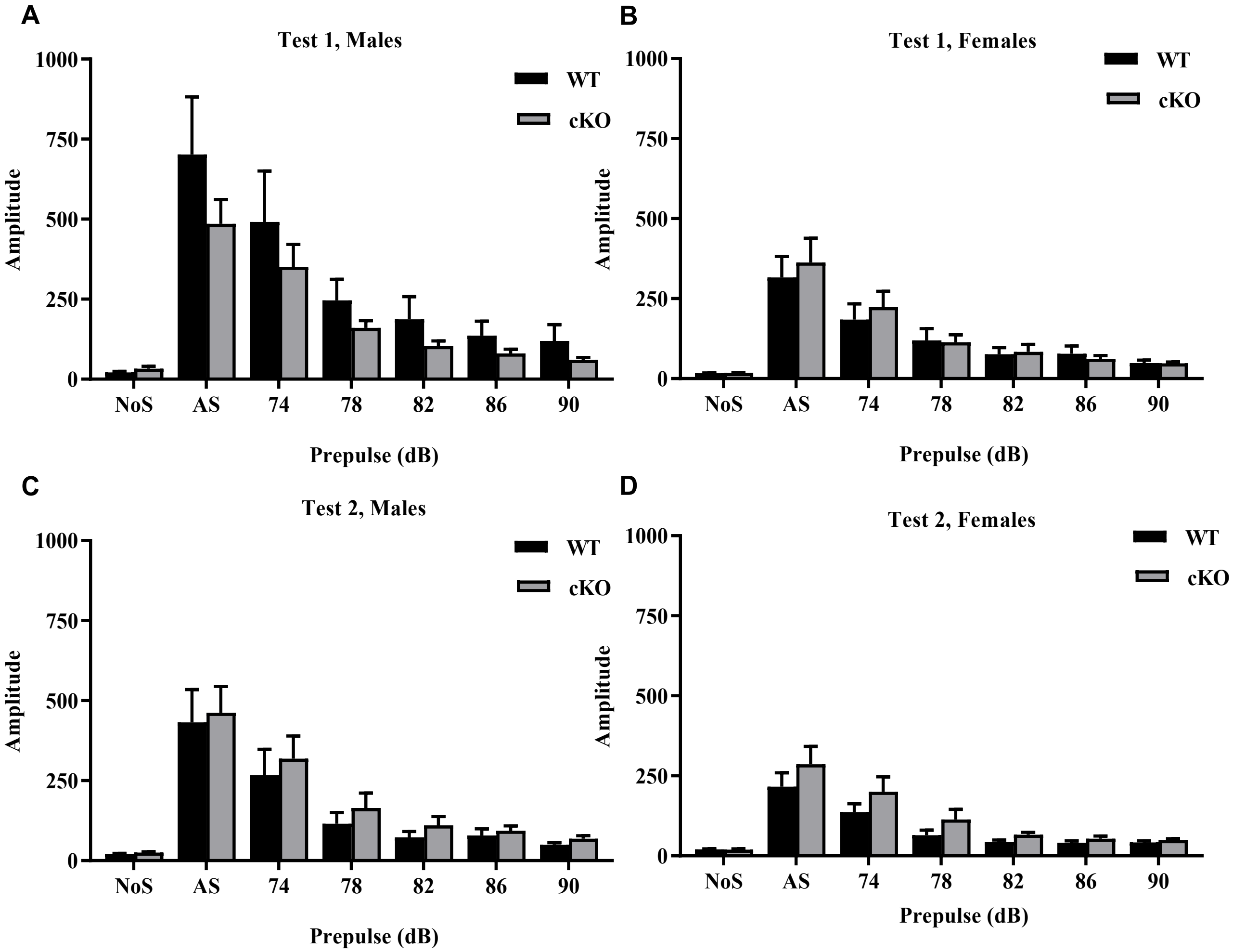

**Supplemental Fig. 4**

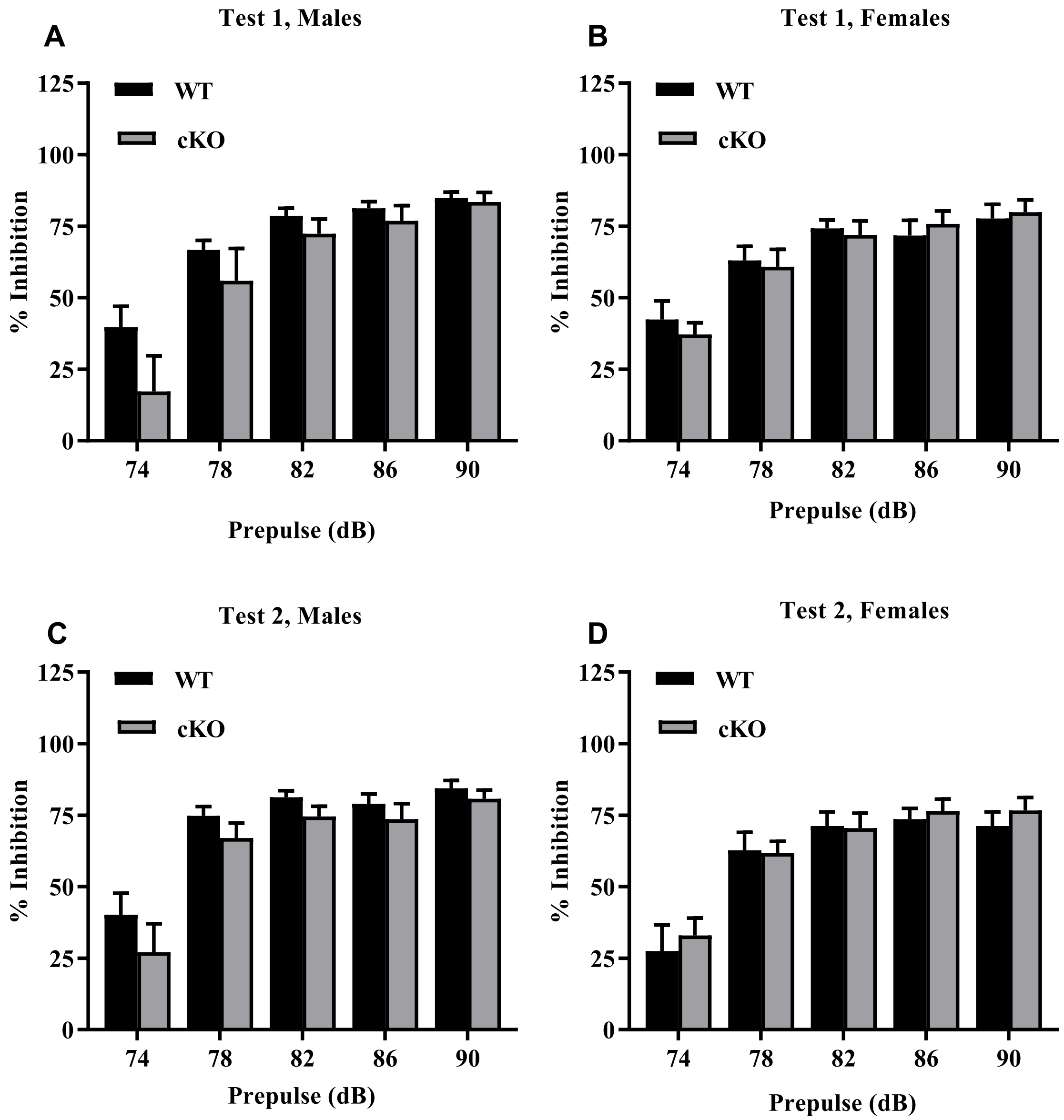

**Supplemental Fig. 5**

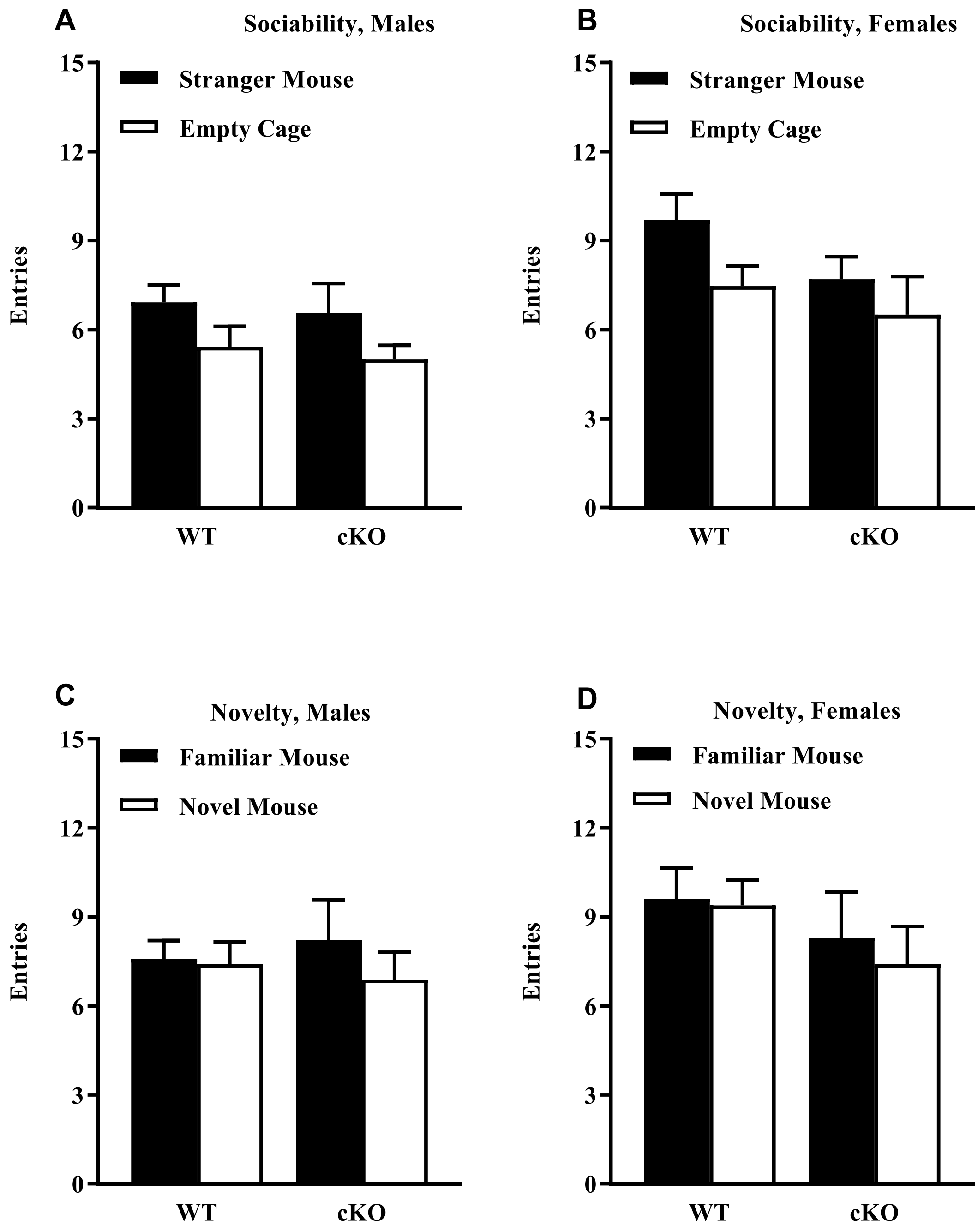

**Supplemental Fig. 6**

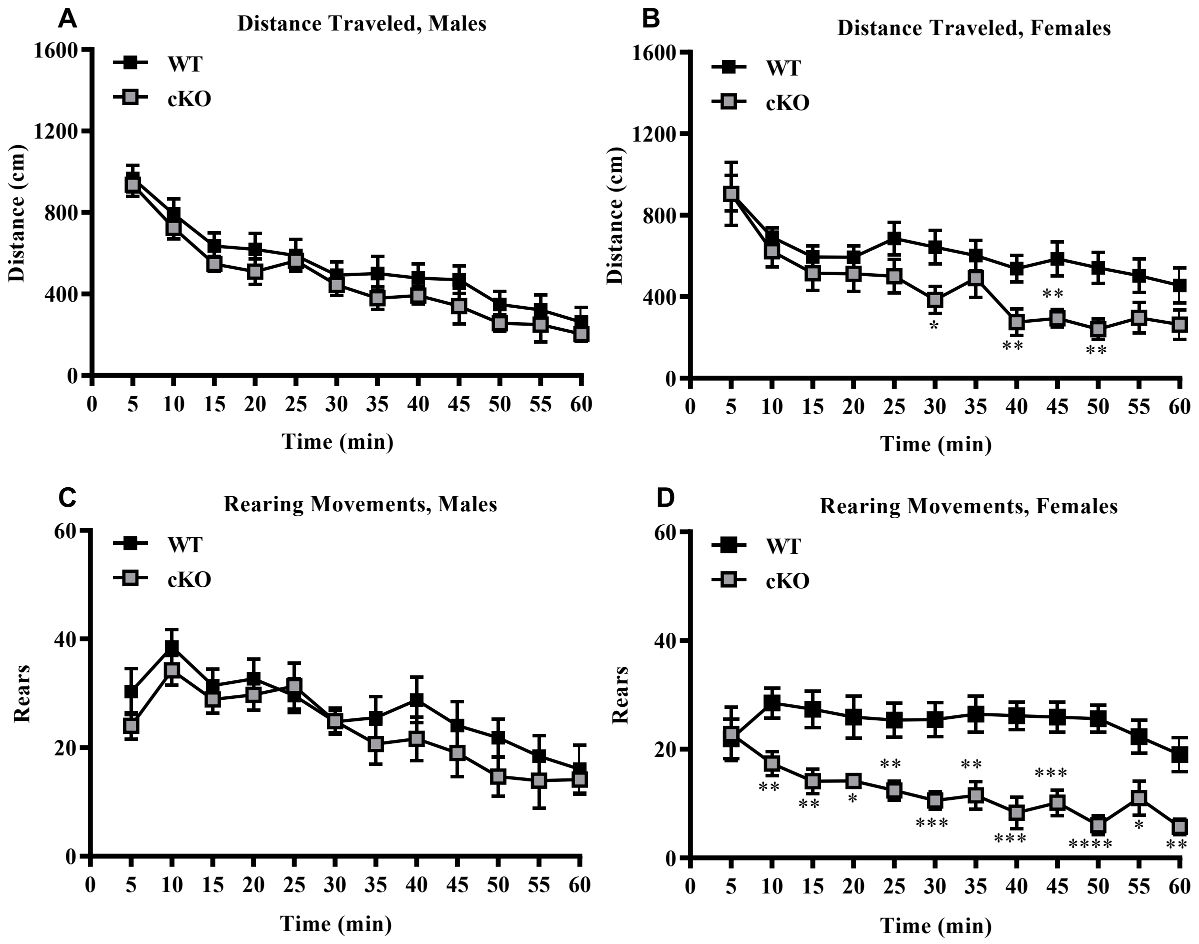

**Supplemental Fig. 7**

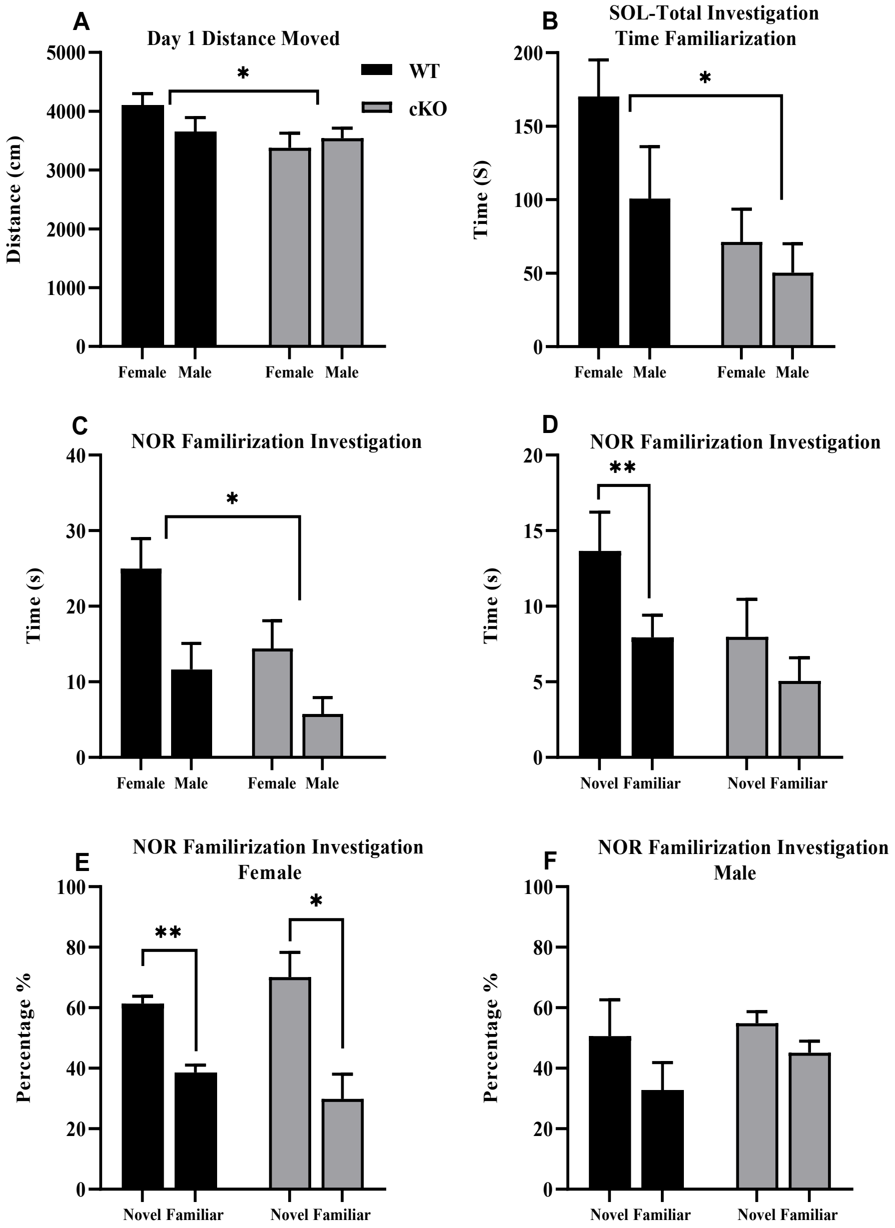

**Supplemental Fig. 8**

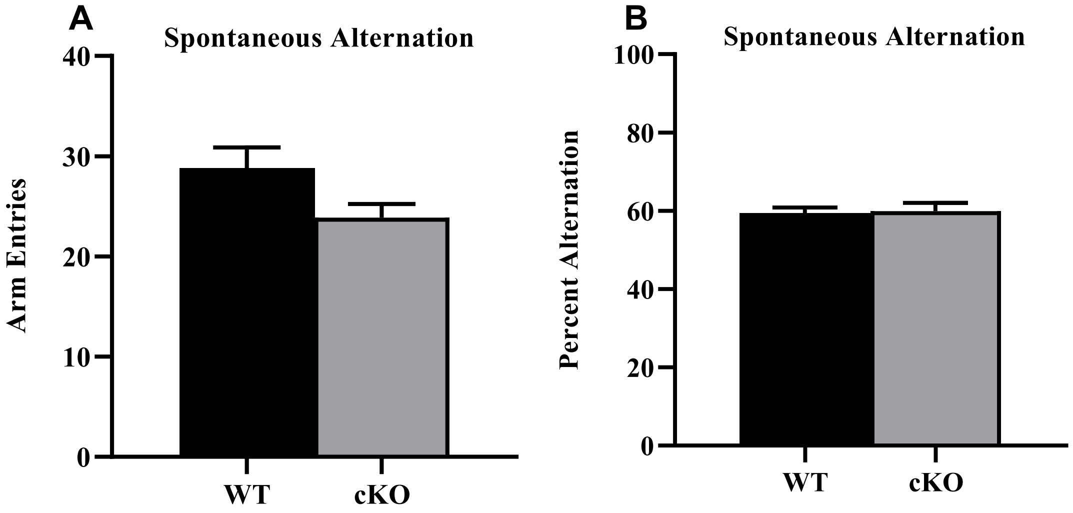

**Supplemental Fig. 9**

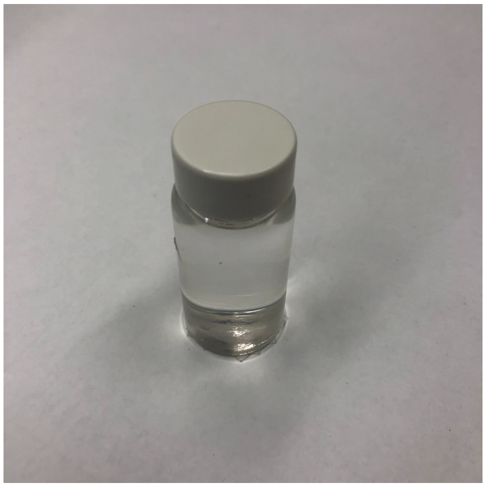

**Object 1 description: Assembled Screw Vial Kit, Clear Glass; 20mL with screw on cap and filled with water.**

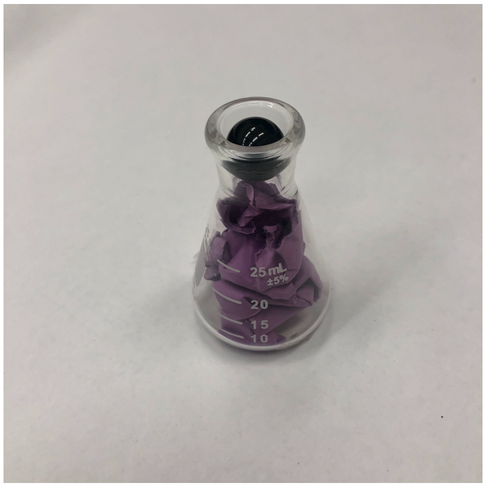


**Object 2 description: Pyrex Glass Erlenmeyer Flask; 25 mL with purple construction paper folded inside and glossy black glass ball insert**

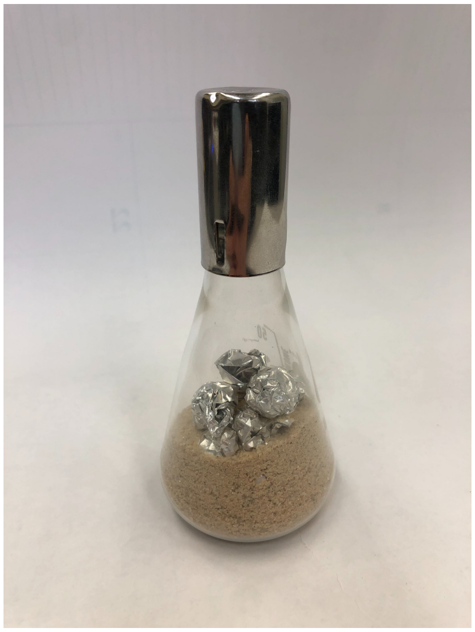

**Object 3 description: Belco 50 mL Erlenmeyer with alumnium foil shavings and sand.**

**Supplemental Fig. 10**

| AGE (WEEKS) | PROCEDURE |
|---|---|
| **6-8** | Elevated plus maze test for anxiety-like behavior. |
| **7-9** | Open field (1 hour). |
| **8-10** | Marble-bury assay. |
| **9-11** | Social approach in a 3-chamber choice task. |
| **10-12** | Rotarod test for motor coordination |
| **11-13** | First acoustic startle test (index of sensorimotor gating). |
| **12-14** | Buried food test for olfactory ability, wire-hang test for grip strength. |
| **14-16** | Morris water maze, visible platform test. |
| **15-17** | Acquisition of spatial learning in the water maze. |
| **16-18** | Reversal learning in the water maze. |
| **17-19** | Second acoustic startle test, hot-plate test for thermal sensitivity. |
